# Termites became the dominant decomposers of the tropics after two diversification pulses

**DOI:** 10.1101/2025.03.25.645184

**Authors:** Simon Hellemans, Menglin Wang, Corentin Jouault, Mauricio M. Rocha, Jaqueline Battilana, Tiago F. Carrijo, Frédéric Legendre, Fabien L. Condamine, Yves Roisin, Eliana M. Cancello, Rudolf H. Scheffrahn, Thomas Bourguignon

**Affiliations:** Okinawa Institute of Science & Technology Graduate University, 1919-1 Tancha, Onna-son, 904-0495 Okinawa, Japan; Essig Museum of Entomology Research, University of California, Berkeley, Berkeley, CA, USA; Oxford University Museum of Natural History, University of Oxford, Parks Road, Oxford OX1 3PW, UK; Museu de Zoologia da Universidade de São Paulo, Avenida Nazaré, 481, Ipiranga, São Paulo/SP, 04263-000 Brazil; Universidade Federal do ABC, Avenida dos Estados, 5001, Santo André, SP, 09210-580 Brazil; Institut de Systématique, Évolution, Biodiversité (UMR 7205), Muséum national d’Histoire naturelle (MNHN), CNRS, Sorbonne Université, EPHE-PSL, Université des Antilles, CP50, 57 Rue Cuvier, 75005, Paris, France; Institut des Sciences de l’Évolution de Montpellier (UMR 5554), Université de Montpellier, CNRS, Place Eugène Bataillon, 34095 Montpellier, France; Evolutionary Biology and Ecology, Université Libre de Bruxelles, 50 avenue F.D. Roosevelt, 1050 Brussels, Belgium; University of Florida, Fort Lauderdale Research and Education Center, 3205 College Avenue, Davie, Florida, 33314, USA

**Keywords:** Kalotermitidae, Mitogenomes, Neoisoptera, Neotropics, Termitidae, Ultraconserved elements

## Abstract

Insects have the highest species richness among animals, but the extent of their diversity and the timing of their diversification remain unclear. Insect diversification is difficult to infer due to the incompleteness of the fossil record. Phylogenetic trees of extant species reconstructed from an exhaustive sampling can be useful to address major evolutionary questions. Here, we investigated the diversification of termites, which comprise 2,995 described species, using estimates of speciation, extinction, and net diversification rates inferred from molecular phylogenies including 2,800 samples representing 1,377 putative species. Termites originated in the Early Cretaceous ∼132 million years ago. Estimated extinction rates were close to zero despite fossil evidence of extinction; therefore, we focused our interpretations on the net diversification rates. Our analyses detected two significant rate shifts. The first shift occurred at the end of the Cretaceous, initially in the Kalotermitidae, then in the Neoisoptera as they started outcompeting Kalotermitidae. The second shift involved multiple lineages of Neoisoptera, especially Termitidae, which diversified as they colonized the world after the global cooling initiated at the Eocene-Oligocene transition. Our results indicate that termites became the dominant insect decomposers of tropical ecosystems as global climate change impacted ecosystems.

## Introduction

The species richness of modern ecosystems results from past diversification events, that is, the sum of speciation events minus extinction events (Schluter & Pennell, 2017). Both of these biological processes drive evolutionary radiations and taxonomic turnover at geological time scales and have long been studied using the fossil record (Knoll & Nowak, 2017). For example, the fossil record provides evidence for five mass extinction events —global and sudden increases in extinction rates over short geological periods— that occurred during the past 450 million years and have been used to establish the geological timescale (Raup & Sepkoski, 1982). Mass extinction events have affected all life forms on Earth, but are most famous for their impact on vertebrates, such as the demise of the non-avian dinosaurs at the end of the Cretaceous, ∼66 million years ago (Ma), which paved the way for the rise of mammals (Simpson, 1937; Rose, 2006; Upham *et al*., 2019; Álvarez-Carretero *et al*., 2022). In contrast, the diversification dynamic of many other groups has been less studied.

The fossil record is often too fragmentary to make inferences about the dynamic of organism diversification (Benson *et al*., 2021). Alternatively, diversification processes can be estimated using time-calibrated phylogenetic trees reconstructed from extant species (Nee, 2006; Ricklefs, 2007; Pennell & Harmon, 2013). For example, time-calibrated trees have been used to investigate the timing of the early diversification of mammals and estimate their speciation and extinction rates by fitting birth-death models (Meredith *et al*., 2011; Stadler, 2011; Álvarez-Carretero *et al*., 2022; Quintero *et al*., 2024). Birth-death models allow estimating the diversification dynamic with its elementary components, birth (speciation) and death (extinction) rates, controlling the rise and fall of species (Nee *et al*., 1994a; b). As phylogenetic trees became increasingly complete and available, these models became more popular and complex to address a wealth of evolutionary questions (Stadler, 2013; Morlon, 2014; Morlon *et al*., 2024). However, these methods have often been criticized (Kubo & Iwasa, 1995; Crisp & Cook, 2009; Quental & Marshall, 2010; Rabosky, 2010). Lately, it has been proposed that some birth-death models are unidentifiable because an infinity of pairs of speciation and extinction rates can equally fit a given phylogenetic tree (Louca & Pennell, 2020), even though timetrees contain information on the tempo and mode of diversification rates (Nee *et al*., 1994b; a). This is especially the case for timetrees reconstructed using exhaustive sampling (Helmstetter *et al*., 2022; Morlon *et al*., 2022), which is rarely achievable for non-vertebrate animals because of their high species richness.

Insects are represented by an estimated 5.5 million extant species (Stork, 2018). Their fossil record is comparatively meager, with less than 50,000 described species, and comprises many temporal, spatial, and taxonomic gaps (Grimaldi & Engel, 2005; Schachat & Labandeira, 2021). Despite its incompleteness, the fossil record has been used to study early insect diversification (Labandeira & Sepkoski, 1993; Condamine *et al*., 2016) and its potential drivers (Jouault *et al*., 2022b, 2024; Peris & Condamine, 2024). Time-calibrated phylogenies have also been used to study the timing of modern insect diversification (Misof *et al*., 2014; Rainford *et al*., 2014), often suggesting that it coincides with the angiosperm terrestrial revolution, the period during which flowering plants became the dominant flora (Benton *et al*., 2022). However, phylogenetic studies have mainly focused on estimating the divergence times between lineages rather than their diversification rates (Espeland *et al*., 2018; Kawahara *et al*., 2019; McKenna *et al*., 2019; Frandsen *et al*., 2024). Indeed, estimating diversification rates from under-sampled phylogenetic trees has been proven difficult (Cusimano & Renner, 2010; Chang *et al*., 2020). Densely sampled species-level phylogenetic trees have been reconstructed for charismatic and moderately diverse clades such as mammals (Upham *et al*., 2019), birds (Jetz *et al*., 2012), or squamates (Tonini *et al*., 2016); but such trees remain out of reach for insects, which contain many hyperdiverse lineages that are difficult to sample exhaustively (Chesters, 2025), such as Coleoptera, Lepidoptera, or Hymenoptera. While impossible for these hyperdiverse lineages, inferring comprehensive timetrees can be achieved for insect lineages comprising a few thousand species like mammals and birds, such as termites.

Termites are a lineage of social cockroaches comprising ∼3,000 described extant species (Krishna *et al*., 2013; Constantino, 2018). They play a key role in modern ecosystems as ecosystem engineers (Jouquet *et al*., 2016), making up 10-20% of the animal biomass in tropical ecosystems (Eggleton *et al*., 1996; Ellwood & Foster, 2004; Rosenberg *et al*., 2023). Consuming vegetal matter at various decomposition stages, from sound wood to humus and soil (Donovan *et al*., 2001; Eggleton & Tayasu, 2001; Bourguignon *et al*., 2011), they represent the primary non-microbial decomposers (Griffiths *et al*., 2019). In addition, termites are the oldest lineage of social insects, with a fossil record dating back to the Early Cretaceous, ∼130 Ma (Thorne *et al*., 2000; Engel *et al*., 2016; Jouault *et al*., 2021), containing ∼217 described fossil species (as of December 2024, (Constantino, 2024). The fossil record of termites shows temporal gaps across all main lineages, which precludes its use to study the termite diversification dynamics. As a result, a few studies have investigated the diversification dynamics of termites using dated phylogenies and have suggested that termite eusociality, as well as complex hindgut structure associated with prokaryotic microbiota, are associated with increased diversification rates as compared to other groups within the superorder Dictyoptera (Davis *et al*., 2009; Legendre & Condamine, 2018; Condamine *et al*., 2020). Other studies have attempted to reveal broader patterns of termite diversification using a phylogenetic backbone of only 402 species, to which the remaining 2,567 species were included using taxonomic data combined with a stochastic polytomy resolution approach (Chang *et al*., 2020; Pie *et al*., 2021). However, no study has yet investigated the diversification dynamic within termites, which requires robust dated phylogenetic trees encompassing a large fraction of termite species diversity.

Here, we aim at determining how and when termites became the dominant decomposers in tropical ecosystems. We reconstruct the most complete time-calibrated phylogenetic trees of termites to study their diversification dynamics over time and across clades. Molecular data of termites have been accumulated at a fast pace. Early termite phylogenetic datasets consisted of a few genetic markers obtained with Sanger sequencing (e.g., (Inward *et al*., 2007; Legendre *et al*., 2008, 2015), but recent datasets include complete mitochondrial genomes (e.g., (Bourguignon *et al*., 2015, 2017; Wang *et al*., 2023), transcriptomes (Bucek *et al*., 2019), and ultraconserved elements (UCEs) (Hellemans *et al*., 2022c, 2024a). These datasets can be used to infer the termite phylogeny and study their diversification dynamic, albeit with a need to improve the species diversity sampling. Thus, we supplemented the previously published mitochondrial genome and UCE data of 1,205 samples with 1,596 newly sequenced samples to infer a time-calibrated tree including >1,300 species (46% of the total species richness). We used the newly reconstructed trees to investigate termite diversification and its likely drivers with birth-death models.

## Results and Discussion

### Reconstruction of a robust termite phylogenetic tree exhaustive for the Americas

Our sampling was representative of the taxonomic diversity of termites, including each extant family and subfamily as defined by Hellemans *et al*. (2024a) and 218 of the 321 currently recognized genera (Figure 1; Supplementary Data 1; Hellemans *et al*., 2024a: Supplementary Note 4). It also included samples from all continents where termites are known to occur, and was especially exhaustive and nearly complete for the Americas (Supplementary Data 1). We reconstructed phylogenetic trees using a maximum-likelihood approach with four datasets: (i) mitochondrial genomes (hereafter: 3CP+; alignment of 18,099 nt), (ii) mitochondrial genomes without third codon positions (3CP-; 14,193 nt), (iii) UCEs combined with mitochondrial genomes (UCEs_3CP+; 36,600 nt), and (iv) UCEs combined with mitochondrial genomes without third codon positions (UCEs_3CP-; 32,694 nt). Herein, we provide a total of 1596 newly sequenced mitogenomes, thereby doubling the 1,205 mitogenomes previously available.

**Figure 1.**
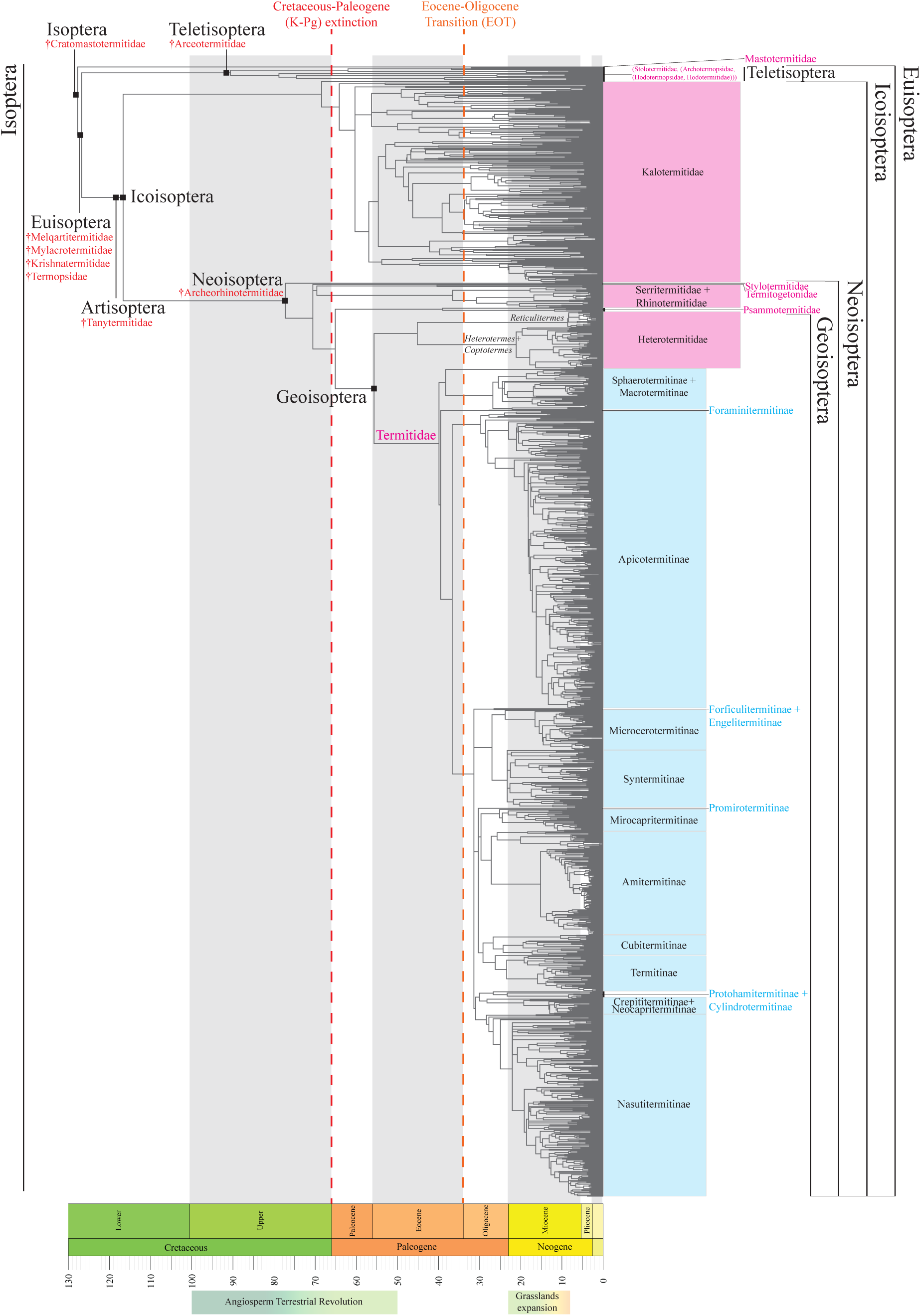
Annotated Termite Tree of Life (TTL) reconstructed using UCEs and mitochondrial genomes without third codon position sites (3CP-). Tips were pruned into 1,377 Operational Taxonomic Units (OTUs) at the 3%-dissimilarity threshold as a proxy for species (for the unpruned tree, see Dryad: File 2). Clades are anchored in the tree by black squares, and extinct families are indicated with the dagger “†” symbol under the clade to which they belong. Family- level ranks are highlighted in magenta, while subfamilies of Termitidae are highlighted in cyan.

We centralize all published mitogenomes in a new curated database repository (available at: https://github.com/oist/TER-MT-DB/) to facilitate future efforts in taxonomy and phylogenetics. Similarly, we enhance the termite UCE database (available at: https://github.com/oist/TER-UCE-DB/) with data from 1,723 new samples (Dryad: File 1), representing more than a five-fold increase from the 315 samples previously available (Hellemans *et al*., 2022c).

Our phylogenetic trees included a total of 2,800 termite samples, with many species represented by multiple samples. We defined Operational Taxonomic Units (OTUs) to approximate species for macroevolutionary analyses. Termite samples were considered conspecific when their mitogenome sequence identity exceeded 97%. While 2 to 4% dissimilarity is generally used to define interspecies boundaries, some specific taxa are considered conspecific with up to 10% dissimilarity (Klimov *et al*., 2019). A 3%-dissimilarity threshold to define species is standard and facilitates downstream investigation of past diversification events. Our four final timetrees comprised 1,377 OTUs, which potentially covers 46% of the 2,993 termite species described worldwide (Krishna *et al*., 2013), including 1,965 samples from the Americas, representing 854 OTUs for 648 species described from the Neotropical and Nearctic realms (Constantino, 2018). These results suggest that our dataset contains many species new to science.

Our dataset was centered on the Americas. For instance, the American Apicotermitinae were represented by 301 OTUs, while the African Apicotermitinae were only represented by 60 OTUs (Supplementary Data 1). These numbers contrast with their reported diversity: 202 species of Apicotermitinae were inventoried in the catalog “*Treatise on the Isoptera of the World*” (Krishna *et al*., 2013), including 112 species from the Afrotropical realm and 38 from the Americas. Therefore, our 301 OTUs of American Apicotermitinae indicate that this group contains at least eight times more species than currently described in the Americas. The actual diversity of this group is likely similar in Africa. We accounted for the discrepancy between sampling efforts and actual species diversity by specifying sampling fractions. However, the diversification pulses identified in this study likely reflect more heavily the history of American termites, especially in the Neotropics, the region hosting the second largest termite diversity after Africa.

Overall, our four phylogenetic trees were largely congruent, as indicated by pairwise comparisons with the generalized Robinson-Foulds method (Smith, 2020), which yielded values ranging from 0.91 to 0.96 after normalization (Supplementary Data 2) —values of one indicating identical tree topologies. The phylogenetic positions of termite families and subfamilies were also largely in agreement with previous termite phylogenetic trees based on mitochondrial genomes (Bourguignon *et al*., 2015, 2017; Wang *et al*., 2023), transcriptomes (Bucek *et al*., 2019), and UCEs (Hellemans *et al*., 2022c, 2024a) —albeit with some topological differences for a few nodes that already presented disagreements among previous phylogenies. Our four timetrees estimated the age of modern termites at 127.0 to 132.0 Ma, which is within the confidence intervals of previous timetrees based on transcriptomes (Buček *et al*., 2019). Reconstructed time divergences were sensibly younger in trees incorporating UCEs than in trees reconstructed solely from mitogenomes (Dryad: File 2), which is in line with a previous study on Kalotermitidae (Buček *et al*., 2022b).

### The estimations of extinction rates are unrealistically close to zero

We used our four termite species trees to investigate the diversification of termites with macroevolutionary models of species diversification. We estimated the variations in rates of speciation (*λ*), extinction (*μ*), and net diversification (*r*) over time with three methods: the Bayesian Analysis of Macroevolutionary Mixtures (BAMM) (Rabosky, 2014; Rabosky *et al*., 2014b); the episodic birth-death model of TreePar (Stadler, 2011); and RPANDA (Morlon *et al*., 2016), which compared six birth-death models with various speciation and extinction scenarios. We ran all analyses by specifying a global sampling fraction of 0.39 to address the incompleteness of our sampling. BAMM analyses were run twice: once using the global sampling fraction only and once using a custom sampling fraction file considering clade-specific sampling fractions (see Methods).

All BAMM analyses yielded extinction rates close to zero along the entire evolutionary history of termites (Figures 2C, E, G), except for the analyses performed on 3CP+ (Figure 2A), in which high *μ* rates were estimated at root age (132 Ma), slowly decreasing until reaching zero ∼65 Ma. Unlike BAMM, TreePar estimated high values of both *λ* and *μ* (Figures 2B, D, F, and H; Supplementary Data 3). Finally, the best-fit birth-death models identified by RPANDA were either (i) a pure-birth constant rate model (BCST) with *λ* = 0.0799 for 3CP+; (ii) a speciation varying through time without extinction model (BTimeVar) with *λ* = 0.1392 for 3CP- and *λ* = 0.1465 for UCEs_3CP-; or (iii) a speciation and extinction varying through time model (BtimeVarDTimeVar) with *λ* = 0.0919 and *μ* = 0.0026 for UCEs_3CP+ (Supplementary Data 4). Therefore, our analyses consistently estimated high and variable speciation rates but often low extinction rates, close to zero for all termites. This concurs with previous studies performed on the Australian Amitermitinae (Heimburger *et al*., 2022), for which model tests evidenced a better fit of pure-birth (Yule) models than birth-death processes. Overall, our analyses support a scenario where termite diversification was essentially the product of speciation with a negligible impact of extinction.

**Figure 2.**
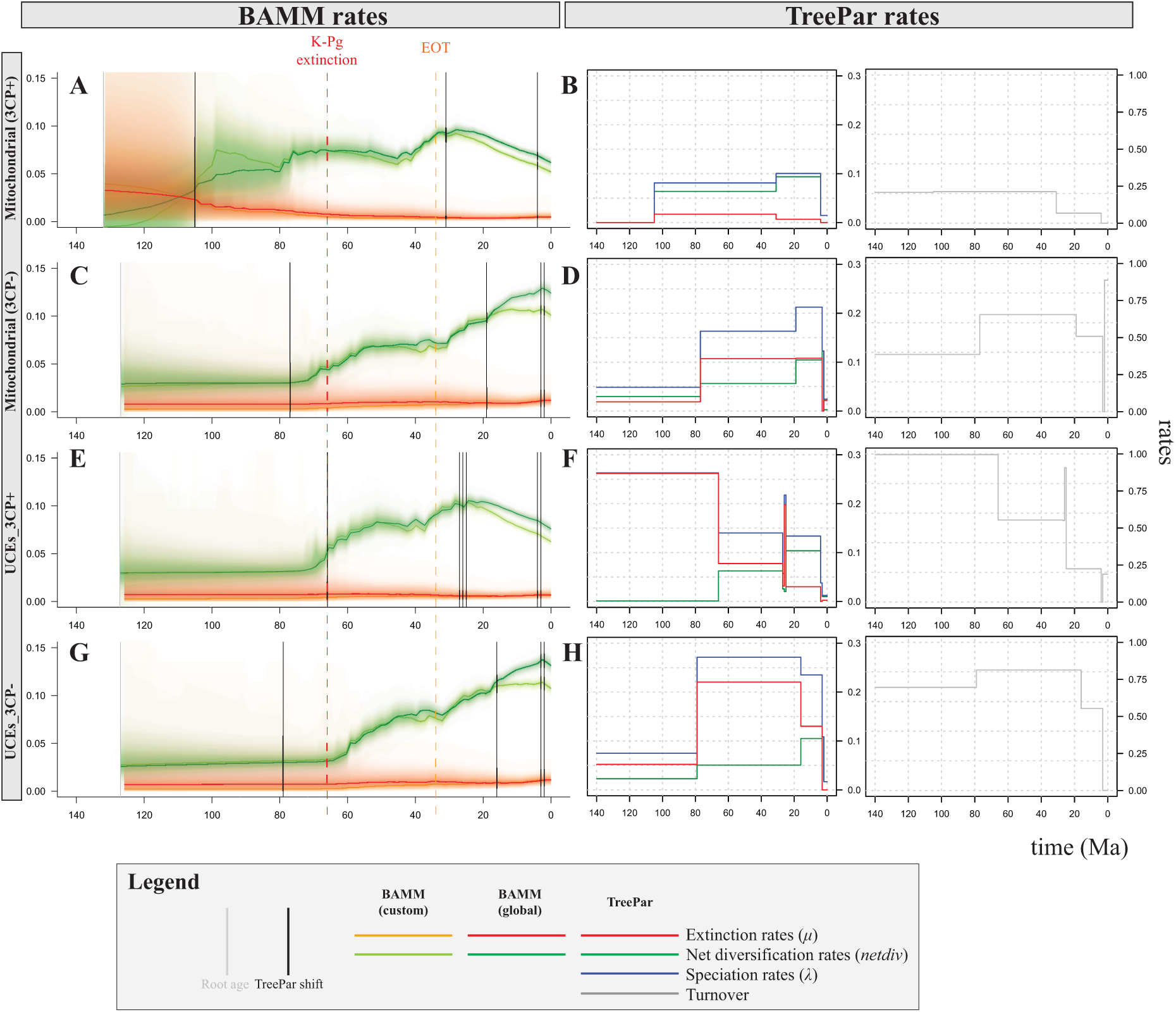
Rates of evolution (extinction, speciation, net diversification and turnover) estimated with BAMM (A, C, E, and G) and TreePar (B, D, F, and H) for four termite phylogenetic trees reconstructed using: (A-B) mitochondrial genomes with third codon positions (3CP+); (C-D) mitochondrial genomes without third codon positions (3CP-); (E-F) UCEs and mitochondrial genomes with third codon position sites (UCEs_3CP+); (G-H) UCEs and mitochondrial genomes without third codon positions (UCEs_3CP-). Vertical bars in BAMM plots represent episodic diversification shifts identified in TreePar’s best model (Supplementary Data 3).

The near absence of extinction during termite evolutionary history suggested herein conflicts with evidence from the fossil record. Termites were already diverse during the Cretaceous —when they were represented by extant but species-poor families such as Mastotermitidae (Jiang *et al*., 2024), Hodotermopsidae (Jouault *et al*., 2022a; Engel & Jouault, 2024), and Stolotermitidae (Zhao *et al*., 2020), and by many species assigned to now extinct families, such as Cratomastotermitidae (Bechly, 2007), Tanytermitidae (Jiang *et al*., 2021), and Archeorhinotermitidae (Krishna & Grimaldi, 2003). The termite fossil record also documents more recent extinction events. For example, fossils of *Mastotermes* from the Miocene have been found in Europe (Emerson, 1965), Africa (Engel *et al*., 2016), and Central America (Krishna & Grimaldi, 1991), while the genus is now restricted to Australia, portraying multiple extinction events across the globe. Overall, the fossil record indicates that termites have experienced extinctions throughout their entire evolutionary history (Engel *et al*., 2009), faulting the scenario of no extinction suggested by most of our diversification analyses.

The low extinction rates estimated by our analyses of species diversification are likely the result of methodological constraints and limitations. Phylogenetic software generally imposes *μ* ≥ 0. Consequently, when estimations of *μ* converge to negative values, which is biologically irrelevant but mathematically possible, extinction rates are estimated as zero by default (Louca & Pennell, 2021). We suspect that the low values of *μ* estimated in this study are the result of these methodological limitations. Because of our unrealistic estimation of a zero-extinction rate, together with the problem of identifiability of *λ* and *μ* —characterized by an infinite number of models with different rates being equally likely (Louca & Pennell, 2020)— we will not consider *λ* and *μ* further in this paper. Instead, we will consider diversification rates, which are less affected by these problems.

### The diversification rates of termites increased in two pulses

Our analyses of termite diversification indicate important fluctuations in diversification rates during their evolutionary history. Our BAMM analyses identified between three and nine independent best diversification rate shifts: three to eight shifts when a global sampling fraction only was specified (Figures 3A, C, E, and G) and five to nine shifts when clade-specific custom sampling fractions were specified (Figures 3B, D, F, and H), indicating that the number of detected shifts was more influenced by the phylogenetic reconstruction method than by the sampling fraction scheme. Notably, analyses with a global sampling fraction only provide an American perspective of termite diversification owing to the focus of this study on American species, while specifying clade-specific custom sampling fractions allows to investigate worldwide diversification shifts. This is well illustrated by the Neotropical Apicotermitinae, which were only found to exhibit a diversification rate shift in analyses with a sampling fraction only (Figures 3C, E, G; diversification shift *l*); and the Macrotermitinae, a subfamily not distributed in America and only exhibiting a rate shift in analyses using clade-specific custom sampling fractions (Figures 3D, H; shift *k*).

**Figure 3.**
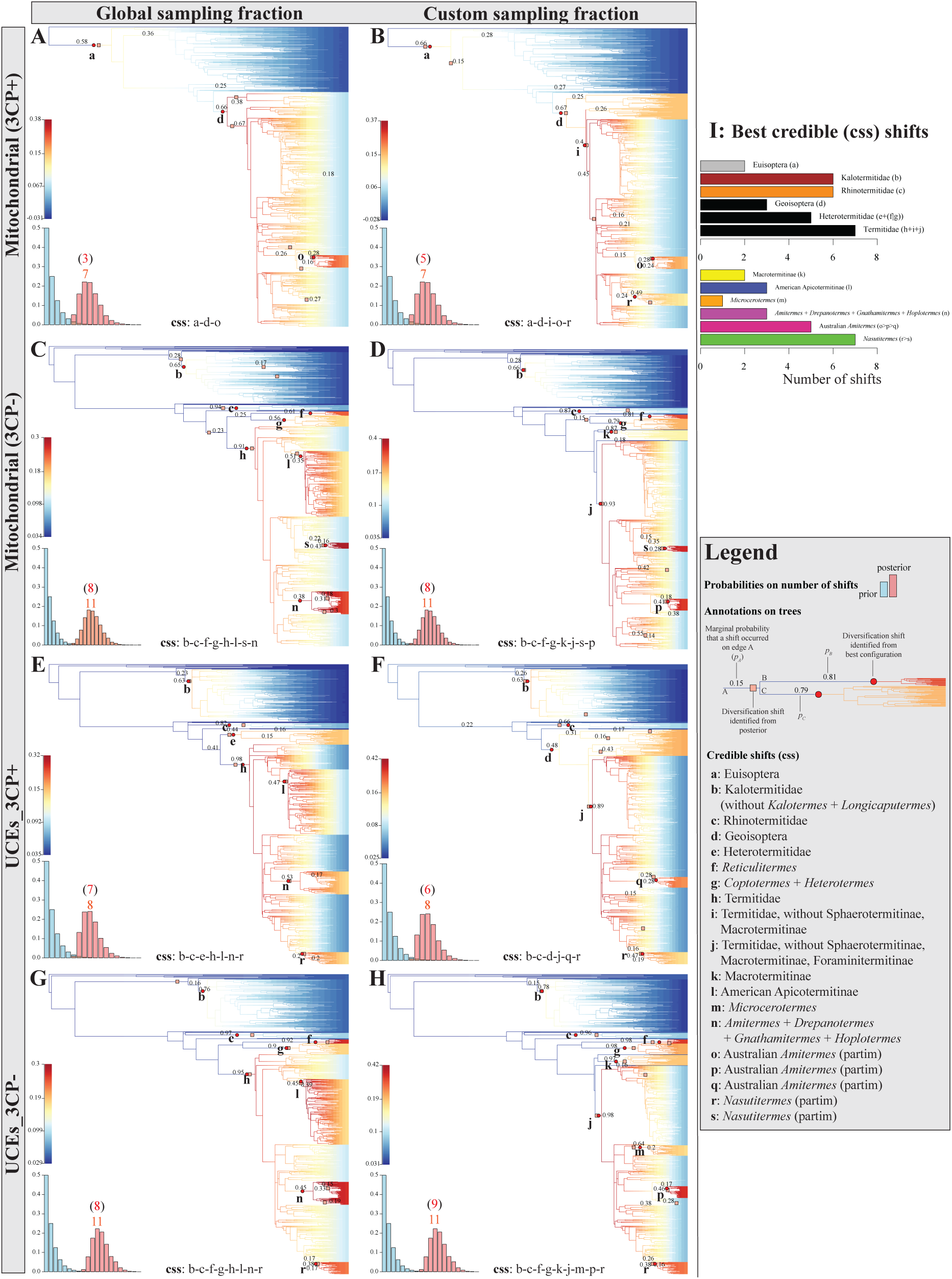
Diversification shifts in Isoptera identified with BAMM for four termite phylogenetic trees reconstructed using: (A-B) mitochondrial genomes with third codon positions (3CP+); (C- D) mitochondrial genomes without third codon positions (3CP-); (E-F) UCEs and mitochondrial genomes with third codon positions (UCEs_3CP+); (G-H) UCEs and mitochondrial genomes without third codon positions (UCEs_3CP-). The analyses were performed using a global sampling fraction of 0.39 (A, C, E, and G), and clade-specific sampling fractions (B, D, F, and H). Barplots represent the prior (blue) and posterior (pink) distribution of shifts. Pink squares are posterior diversification shifts, while red circles are the best credible shifts identified by BAMMtools’ ‘credibleShiftSet’ function (“css” shifts; e.g. seven shifts were identified from the posteriors, three being in the credible set in subpanel A). The trees’ branch colors represent net diversification rates. Branch labels indicate the marginal probability (*p*; indicated when *p* ≥ 0.15) with which a shift occurred on that edge. (I) Summary of the credible css shifts over the eight analyses. Shifts assigned to clades below the subfamily rank (n-s) have their members listed in Supplementary Data 7.

The branches on which the best shifts are located varied among analyses, in part because BAMM identifies areas in trees where shifts are likely to occur rather than precise locations (Rabosky *et al*., 2014a; b). All best shifts represented increases in diversification rates. The two types of analyses, performed with a global sampling fraction only and with custom sampling fractions, identified between 30% (three shifts over 10 for UCEs_3CP+; Figures 3E-F) and 60% of best shifts on the same branches (three shifts over five for 3CP+; Figures 3A-B). One best shift was found outside the Neoisoptera, either at the origin of Kalotermitidae (Figures 3C-H; shift *b*) or on the branch leading to Euisoptera (Figures 3A-B; shift *a*), which includes all termites except for Mastotermitidae (Figure 1). All other best shifts were located within the Neoisoptera. Some general trends can be summarized (Figure 3I): (i) increase in diversification rates occurred early on in the evolution of Neoisoptera, represented either by one shift at the base of the Geoisoptera (Heterotermitidae and Termitidae; Figures 3A, B, and F; shift *d*) or by multiple shifts around the base of Termitidae, Rhinotermitidae, and Heterotermitidae (Figures 3C-E and G-H; shifts *c, e, h*); (ii) multiple increases in diversification rates occurred within Termitidae, including potential shifts within Apicotermitinae, Microcerotermitinae, Amitermitinae, and Nasutitermitinae (Figures 3A-H).

The BAMM results essentially concur with the TreePar analyses, which identified two main increases in diversification rates during the termite evolution, with timing varying among analyses (Figure 2). The first increase in rate occurred during the Late Cretaceous and was dated between 105 to 66 Ma (Supplementary Data 3), corresponding to the shift identified by BAMM near the origin of Kalotermitidae (Figure 3; shift *b*). This shift also reflects the early burst among Neoisoptera (Figure 3; shifts *c*-*e*, *h*), which compensated for the decreasing diversification rate of Kalotermitidae early on in the Paleogene. The second increase in rates occurred during the Oligocene-late-Miocene period and was dated between 31 and 16 Ma, likely corresponding to the multiple shifts detected by BAMM within the neoisopteran families (Figure 3; shifts *f*, *g*) and, more importantly, within most Termitidae subfamilies (Figure 3; shifts *i*-*s*). Overall, our results highlight two main diversification pulses that gave rise to modern termite diversity (Figure 4). While our sampling bias could affect the number of recent shifts, it is unlikely that such sampling would have impacted more ancient shifts.

**Figure 4.**
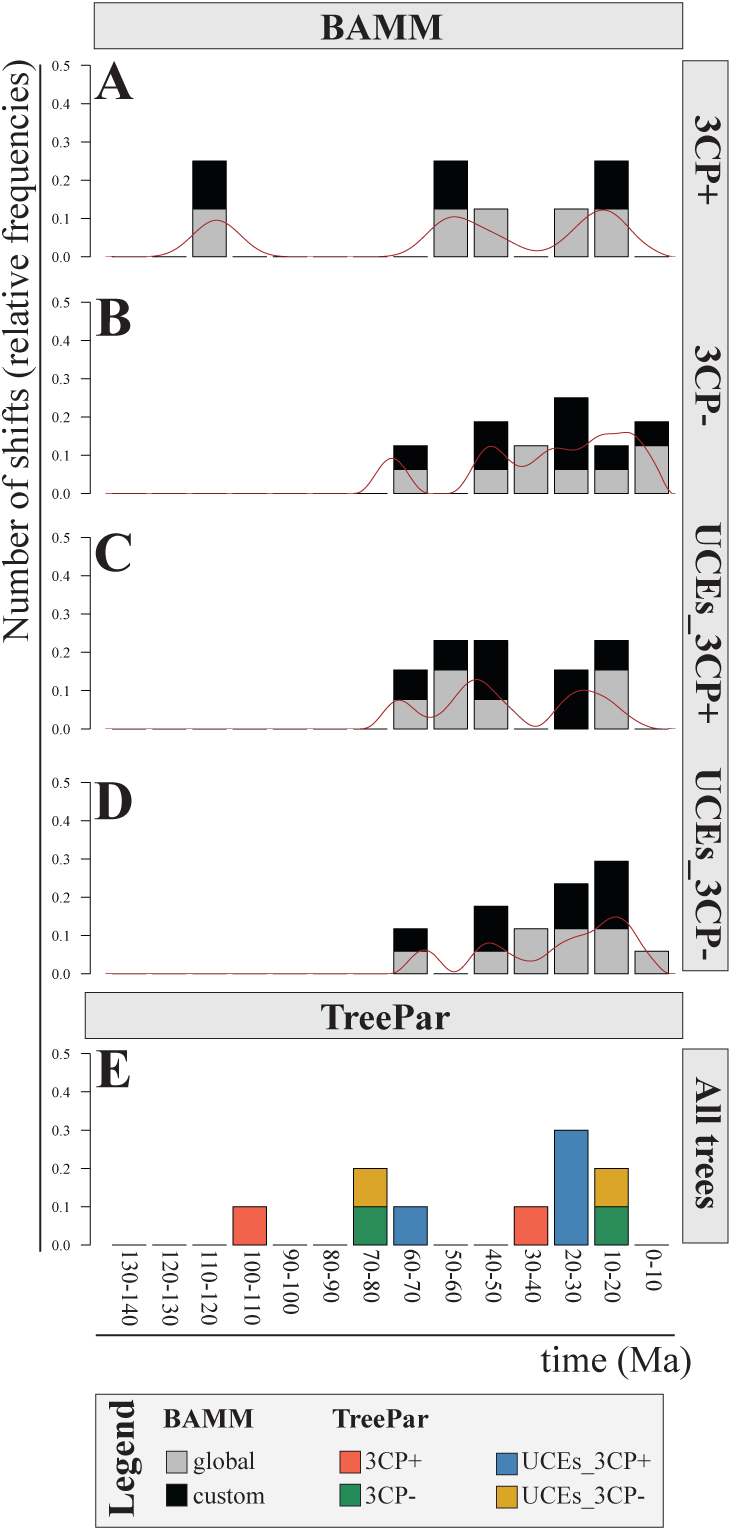
Temporal synthesis of shifts identified by BAMM (A-D) and TreePar (E) for four termite phylogenetic trees (see also Figures 2-3). Shifts frequencies were plotted across time using barplots (with bins spanning 10 Ma) and smoothed density curves. Shifts occurring in the last 5 Ma were not considered in TreePar (see Methods). Binned BAMM shifts retrieved in analyses with either global or clade-specific sampling fractions are represented in grey or black, respectively, while TreePar shifts were synthetized in a single chart with one color per analysis.

### Contrasted diversification patterns within termites: Kalotermitidae slowed down as Neoisoptera diversified

The family Kalotermitidae was among the first termite lineage to experience a diversification rate shift during the Cretaceous/Paleogene boundary, between 69 and 61 Ma (Figures 3C-H; shift *b*). This shift was marked by an increased diversification at the crown of Kalotermitidae, followed by a rapid slowdown coinciding with the first diversification rate shifts within Neoisoptera during the Cenozoic, between 58 and 46 Ma (Figures 3A-H; shifts *c*, *d*). We hypothesize that the diversification slowdown in Kalotermitidae may have been a consequence of the increasing diversification rate in Neoisoptera, reflecting competition for resources between two sister lineages of wood decomposers. Notably, this trend initiated at the end of the Cretaceous is still ongoing nowadays, with Kalotermitidae and Neoisoptera being competitors in modern ecosystems (Chouvenc *et al*., 2021; Buček *et al*., 2022b). Kalotermitids are frequently found in areas where competing wood-feeding Neoisoptera are scarce or absent (e.g., arid regions and remote islands; Buček *et al*., 2022b). In Neoisoptera-rich regions, Kalotermitidae are primarily found in places that wood-feeding Termitidae and other Neoisoptera have difficulties accessing, such as isolated dead branches in the tree canopy (Roisin *et al*., 2006) and mangrove trees (Scheffrahn *et al*., 2006) or the hardwood at the center of tree trunks (personal observation). The competitive superiority of Neoisoptera is also reflected in their diversity, which exceeds that of Kalotermitidae: among the 2,933 extant described termite species listed in Krishna et al. (2013), 456 are Kalotermitidae and 2,439 Neoisoptera, including 2,076 species of Termitidae, one-third to half of which wood feeders (Krishna *et al*., 2013). Mastotermitidae and Teletisoptera (Archotermopsidae, Hodotermitidae, Hodotermopsidae, and Stolotermitidae) may have been outcompeted by Neoisoptera as well. In South America, Mastotermitidae persisted at least until the Miocene (Engel *et al*., 2009), which roughly coincides with the colonization of the Neotropics by the wood-feeding termitids *Nasutitermes* and *Microcerotermes* (Bourguignon *et al*., 2017). Our results suggest that the rise to dominance of Neoisoptera was initiated before the origin of Termitidae, the family including the primary decomposers of organic matter in modern tropical and subtropical terrestrial ecosystems (Eggleton *et al*., 1996; Bignell & Eggleton, 2000).

### The diversification of multiple lineages coincides with their worldwide dispersal after the Eocene-Oligocene transition

Most of the diversification shifts in BAMM occurred between the Oligocene and the middle Miocene and involved multiple lineages within Geoisoptera, which includes Heterotermitidae and Termitidae (Figure 3). These events were captured by TreePar in the second diversification pulse dated between 31 and 16 Ma (Figures 2B, D, F, and H; Figure 4). Notably, these diversification shifts occurred after the Eocene-Oligocene transition (EOT; 34 Ma), a period of global cooling (Ivany *et al*., 2000; Zachos *et al*., 2001; Goldner *et al*., 2014). The EOT likely contributed to the retraction of the once widely distributed megathermal rainforests toward the tropical and equatorial regions (Morley, 2011), leading to the present-day latitudinal diversity gradient. The Miocene also saw the expansion of grass-dominated savanna biomes in response to climatic shifts toward arid conditions, about 23–8 Ma (Jacobs, 2004; Edwards *et al*., 2010; Strömberg, 2011). As important decomposers in both tropical rainforests and savannas (Collins, 1981; Griffiths *et al*., 2019), Geoisoptera diversified and likely filled the available niche of decomposers in ecosystems undergoing major turnovers and susceptible to invasion.

The diversification shifts in Geoisoptera from the early Oligocene to the middle Miocene also coincide with their spread around the world through transoceanic dispersals (Bourguignon *et al*., 2017; Buček *et al*., 2022b; Hellemans *et al*., 2022a; Wang *et al*., 2023). We performed ancestral range estimation using the Dispersal-Extinction-Cladogenesis (DEC) model implemented in BioGeoBEARS (Matzke, 2013) considering six biogeographic realms adapted from Holt *et al*. (2013) (Supplementary Figure 1). We estimated that termites dispersed among biogeographic realms on average 96.42 times over our 100 simulations. Dispersal events during the Cretaceous are likely artefactual because ancient disjunctions in the distribution of early-diverging lineages such as Teletisoptera are due to vicariance (Wang *et al*., 2022). We detected an average of 55.66 dispersal events involving Neoisoptera, 50.32 of which occurred after the EOT, and 41.16 dispersal events involving Geoisoptera, 39.62 of which occurred after the EOT. For comparison, 35.17 dispersal events were detected in the Kalotermitidae, 23.79 of which happened after the EOT, in line with previous studies (Buček *et al*., 2022b).

Our biogeographic and diversification analyses refine existing models explaining the rise of termites as the main non-microbial decomposers of terrestrial tropical ecosystems. The dispersal of Geoisoptera occurred in two waves: the first phase was initiated after the EOT global cooling (34 Ma) as megathermal rainforests retracted to equatorial regions, while the second phase coincided with the expansion of savannas (23–8 Ma; Figures 1,3). Both phases were coupled with increases in diversification rate for Heterotermitidae and multiple lineages of Termitidae, such as within Apicotermitinae, Microcerotermitinae, Amitermitinae, and Nasutitermitinae. Finally, the increased dispersal and diversification rates occurred as Termitidae evolved new feeding strategies, with several lineages specializing in a specific state of organic matter along the wood-soil decomposition gradient (Inward *et al*., 2007; Bourguignon *et al*., 2017; Bucek *et al*., 2019; Chouvenc *et al*., 2021; Hellemans *et al*., 2022a). The rise of termites, especially Termitidae, therefore involved simultaneous increases in species diversification, dispersal, and ecological diversification.

### Termite diversification correlates best with Oxygen and Carbon dioxide concentration

The ∼130-million-year period of the termite evolutionary history has experienced various environmental fluctuations. We investigated the role of past environmental changes on termite diversification using RPANDA. Specifically, we test for a correlation between changes in termite diversification rates and variations of six biotic and abiotic variables: O2 and CO2 atmospheric concentrations, global temperature, continental fragmentation, and relative diversity of angiosperms and gymnosperms. We found that termite diversification rates correlate best with variations in O2 concentrations for three of the four trees (3CP+, UCEs_3CP+, and UCEs_3CP-) and with variations in CO2 concentration for the other tree (3CP-; Supplementary Data 5). The high positive values of *α* and *β* —the coefficients of correlation specific to *λ* and *μ*— ranging from *α* = 5.05 to 10.02 and from *β* = 6.10 to 11.27 indicate strong positive correlations with O2 concentrations, suggesting that both speciation and extinction rates increased as O2 concentration increased. For the fourth tree, we found low positive values of *α* and *β* (*α* = 0.0001; *β* = 0.004) indicative of a weak positive correlation with CO2 concentration, suggesting that both speciation and extinction rates increased as CO2 concentration increased. From the Late Triassic to the present, the O2 concentration rose essentially continuously (Berner, 2006). Similarly, termite diversification has largely continuously increased since the end of the Cretaceous (Figures 2-3), which is congruent with the recovered correlation with O2 concentration. However, the precise nature of the link between termite diversification and fluctuations in O2 concentration should be interpreted with caution as estimations of O2 atmospheric concentration from the Late Triassic to the Cenozoic show large-scale variations depending on modeling approaches (Berner *et al*., 2007; Harrison *et al*., 2010). Importantly, the angiosperm terrestrial revolution, which denotes the rise of angiosperms and the collapse of gymnosperms since the end of the Cretaceous, has been suggested as an important contributor to the diversification of various insects, such as pollinators (Benton *et al*., 2022), was not recovered as significantly contributing to termite diversification. Although it may seem unexpected given the plant-termite association, this result is in line with the observations that termites are generalist in terms of plant taxa they feed on (Bourguignon *et al*., 2011), with few termite species being specialists (Miura & Matsumoto, 1997; Martius *et al*., 2000; Poissonnier *et al*., 2018).

### Conclusions

We studied the diversification of termites using multiple methods and molecular timetrees inferred from a large sampling. Our approach allowed assessing the robustness of popular methods designed to study past diversification events. Many studies using phylogenies of extant species and macroevolutionary models to infer past diversification events rely upon a single tree and methods to construct narratives. Among the most important results to better understand the termite evolutionary history, we found two main diversification periods and we found strong positive correlations between diversification and O2 concentrations, suggesting that both speciation and extinction rates increased as O2 concentration increased. The first net diversification pulse occurred at the end of the Cretaceous, initially in Kalotermitidae and then in Neoisoptera. The second pulse involved multiple lineages of Neoisoptera, especially Termitidae, as they colonized the world after the global cooling initiated at the EOT. In addition to shedding new light on the diversification history of termites, our study provides a wealth of new data, including the mitogenomes and UCEs from 1,596 new samples, thereby more than doubling the amount of available data for these molecular markers. The molecular data has previously been used for taxonomic revisions and new species descriptions in the Amitermitinae (Scheffrahn, 2024), Apicotermitinae (Carrijo *et al*., 2023), and Nasutitermitinae (Scheffrahn *et al*., 2025). However, many species and genera remain to be described from the dataset described here, as evidenced by the presence of 854 species-level OTUs from the Americas in our dataset for 648 termite species described from the regions (Constantino, 2018). To facilitate future efforts in taxonomy and phylogenetics, we centralized all published mitogenomes in a new curated database repository (available at: https://github.com/oist/TER-MT-DB/). We hope that this new database, together with the UCE database (Hellemans *et al*., 2022c; available at: https://github.com/oist/TER-UCE-DB/), will serve as the base for future taxonomic revision of termites.

## Material and Methods

### Biological samples

We gathered a dataset comprising sequence data from 2,800 samples, including 1205 previously published mitochondrial genomes (Cameron & Whiting, 2007; Tokuda *et al*., 2012; Wei *et al*., 2012; Cameron *et al*., 2012; Chen *et al*., 2014, 2021; Bourguignon *et al*., 2015, 2016; Kai *et al*., 2015; Li *et al*., 2015, 2018; Bourguignon *et al*., 2017; Dietrich & Brune, 2016; Meng *et al*., 2016; Su *et al*., 2016; Wang *et al*., 2016, 2019, 2022, 2023; Zhao *et al*., 2016; Han *et al*., 2017; Hervé & Brune, 2017; Lee *et al*., 2017; Liao *et al*., 2018a; b; Scheffrahn *et al*., 2018, 2025; Wu *et al*., 2018; Forni *et al*., 2019; He *et al*., 2019b; a; Liu *et al*., 2019; Ye *et al*., 2019; Romero Arias *et al*., 2021, 2024; Heimburger *et al*., 2022; Hellemans *et al*., 2022c; a, 2025; Buček *et al*., 2022b; Arora *et al*., 2023b; Qian *et al*., 2023; Yu *et al*., 2023; Carrijo *et al*., 2023; Gergonne *et al*., 2024) and sequences from 1596 newly sequenced samples, mostly of American origin (Supplementary Data 1). Samples are stored in the Museu de Zoologia da Universidade de São Paulo (Brazil), the University of Florida Termite Collection (U.S.A.), the Université Libre de Bruxelles (Belgium), and the Okinawa Institute of Science and Technology (Japan).

### DNA sequencing

Whole genomic DNA was extracted using the DNeasy Blood & Tissue extraction kit (Qiagen). Library preparation was performed using the NEBNext® UltraTM II FS DNA Library Preparation Kit (New England Biolabs) and the Unique Dual Indexing Kit (New England Biolabs), with reagent volumes reduced to one-fifteenth of the manufacturer’s recommendations. For samples stored in ethanol for over 20 years, whose DNA is typically fragmented, the incubation time of the enzymatic fragmentation step was set to a maximum of five minutes. Other steps were performed as recommended by the manufacturer. Libraries were pooled in equimolar concentration and paired-end sequenced using the HiSeq 4000, HiSeq X, and NovaSeq 6000 Illumina platforms at a read length of 150 bp.

### Assemblies and annotation of target loci

Raw reads were quality-trimmed using fastp *v*0.20.1 (Chen *et al*., 2018) and assembled with metaSPAdes v3.13 (Nurk *et al*., 2017). Mitochondrial scaffolds were first identified in the metagenome assemblies using MitoFinder *v*1.4 (Allio *et al*., 2020). Fragments from incomplete mitogenomes were elongated using TCSF_IMRA *v*2.7.1 (Kinjo *et al*., 2015); and mitogenomes that failed this step were contiguated using BWA-MEM *v*0.7.10 (Li, 2013) and the consensus function of samtools *v*1.15.1 (Li *et al*., 2009). Mitogenomes were then annotated using MitoFinder. UCEs and their flanking 200 bp at both 5’ and 3’ ends (∼600 bp loci) were extracted from metagenome assemblies using PHYLUCE *v*1.6.6 (Faircloth, 2016), LASTZ (Harris, 2007), and a termite-specific bait set targeting 50,616 loci (Hellemans *et al*., 2022c).

Newly generated mitochondrial genomes are available on GenBank (see accession numbers in Supplementary Data 1). Additionally, we provide a curated termite mitogenome database encompassing all published records, available at: https://github.com/oist/TER-MT-DB/. The UCE dataset produced in this study is available on the Dryad Digital Repository (File 1), available at: XXX. This paper represents the contribution #5 to the Termite UCE Database, available at: https://github.com/oist/TER-UCE-DB/.

### Sequence alignments

Each gene of the 2,800 mitogenomes included in this study was aligned independently. The two mitochondrial rRNA genes and the 22 mitochondrial tRNA genes were aligned as DNA sequences using MAFFT *v*7.305 (Katoh & Standley, 2013). For each of the 13 mitochondrial protein-coding genes, DNA sequences were translated into protein sequences with the transeq function from EMBOSS *v*6.6.0 (Rice *et al*., 2000) and aligned with MAFFT. Protein alignments were back translated into codon alignments using PAL2NAL *v*14 (Suyama *et al*., 2006). All 37 genes were concatenated using FASconCAT-G_v1.04.pl (Kück & Longo, 2014). The resulting supermatrix was separated into five partitions: combined rRNAs, combined tRNAs, and combined first, second, and third codon positions of protein-coding genes.

UCE data were available for 1,969 of the 2,800 samples considered in this study. Each UCE was aligned with MAFFT implemented in phyluce_align_seqcap_align. Alignments were trimmed internally using Gblocks implemented in phyluce_align_get_gblocks_trimmed_alignments_from_untrimmed (Castresana, 2000; Talavera & Castresana, 2007) with default parameters. Internally trimmed alignments including less than 57% of all samples were filtered out using phyluce_align_get_only_loci_with_min_taxa. In total, 94 loci with sample completeness higher than 57% were retained and combined in a single supermatrix using phyluce_align_format_nexus_files_for_raxml. The size of our UCE supermatrix was 18,501 nucleotides, which is comparable to the ∼15 kilobase pairs of termite mitogenomes. We used a high percentage of completeness for loci selection in order to reduce the size of our supermatrix below the limits imposed by the available computational resources, allowing the completion of downstream phylogenetic analyses. For similar reasons, UCE loci were combined in a single partition for all phylogenetic reconstructions.

### Phylogenetic analyses and molecular dating

We reconstructed four phylogenetic trees using combinations of UCE and mitochondrial genome alignments with and without third codon positions. The four datasets used for phylogenetic reconstructions consisted of (i) the mitochondrial genome alignment with third codon positions included (hereafter: 3CP+), (ii) the mitochondrial genome alignment without third codon positions (3CP-), (iii) the UCE alignment combined with the 3CP+ mitochondrial genome alignment (UCEs_3CP+), and (iv) the UCE alignment combined with the 3CP- mitochondrial genome alignment (UCEs_3CP-).

All trees were reconstructed using the maximum-likelihood approach implemented in IQ-TREE v2.2.2.5 (Minh *et al*., 2020) with a phylogenetic terrace-aware data structure (Chernomor *et al*., 2016). To assess branch supports, we ran 1,000 replicates with both the ultrafast bootstrap (UFB) algorithm and the Shimodaira–Hasegawa-like approximate likelihood-ratio test (SH-aLRT) (Guindon *et al*., 2010; Hoang *et al*., 2018). No model selection test was performed, and the GTR+F+I+G4 model was set for all partitions. This approach improved the computational feasibility of this study. The GTR+F+I+G4 model was chosen as it generally produces similar inferences to the best-fit nucleotide substitution model (Abadi *et al*., 2019). In addition, this model was selected as the best-fit model in previous termite phylogenetic studies using mitogenomes (Hellemans *et al*., 2022a). The topologies of the trees were constrained to ensure the monophyly of Macrotermitinae + Sphaerotermitinae and non- Macrotermitinae non-Sphaerotermitinae Termitidae, two lineages proven to be monophyletic by phylogenetic reconstructions performed on transcriptome and UCE data (Bucek *et al*., 2019; Hellemans *et al*., 2022c, 2024a) but branching differently in mitogenome-based phylogenetic trees (Hellemans *et al*., 2017; Wang *et al*., 2023). Reconstructed trees were dated with the least square dating method using LSD2 (To *et al*., 2016) implemented in IQ-TREE. Node dating was performed using time intervals compatible with fossil data and previous time-calibrated Bayesian phylogenetic reconstructions (Hellemans *et al*., 2022a; Supplementary Data 6) with the options “--date-ci 100 --date-tip 0”.

Our phylogenetic trees were reconstructed using 2,800 samples, many of which belonged to the same termite species. We simplified our phylogenetic tree to a species-level tree for downstream analyses. We used operational taxonomic units (OTUs) comprising samples exhibiting over 97% similarity over the entire length of their mitogenome as a proxy for termite species. This classification approach resolves the problems linked to species misidentification. Samples were clustered into OTU using all-versus-all blast with megaBLASTn v2.13.0+ (Camacho *et al*., 2009). The 2,800 samples used in this study were clustered into 1,377 OTUs. Dated trees were pruned to keep one sample per OTU and no outgroups using the “drop.tip” function of the R package ‘ape’ (Paradis *et al*., 2004). Tree topologies were compared using the normalized Generalized Robinson-Foulds tree comparison metrics with the ’NyeSimilarity’ function of the R package ‘TreeDist’. We used Mesquite v3.7 (Maddison & Maddison, 2019) to resolve polytomic nodes. Zero-length branches were given a length of 0.0000001 for downstream diversification analyses. All trees are available from the Dryad repository (File 2).

### Estimation of diversification rates through time

We performed the diversification analyses using all four dated OTU trees. We used BAMM v2.5 and its associated R package BAMMtools v2.1.11 (Rabosky, 2014; Rabosky *et al*., 2014b) to estimate the rates of speciation (*λ*), extinction (*μ*), and net diversification (*r*) through time and identify the placement of potential shifts on the phylogenetic tree. BAMM models the evolutionary dynamics of lineages (macroevolutionary cohorts) through time. Cohorts share *λ* and *μ* rates, thence they also share net diversification *r* rates, and are delimited by diversification rate shifts (Rabosky *et al*., 2013, 2014a). The diversification analyses were performed on the largest termite phylogenetic trees of life assembled to date, which included 1,377 species-level OTUs. We considered the number of extant termite species to be 3,500, which roughly corresponds to the number of all described species, including synonyms (see Constantino, 2018). Therefore, our sampling corresponds to 39% of the described termite fauna.

We addressed the possible issues linked to the heterogeneity of our sampling effort by assigning a specific sampling fraction to genera of Mastotermitidae and Teletisoptera, as well as each family of Icoisoptera (Kalotermitidae, and non-Termitidae Neoisoptera), and each subfamily of Termitidae (Dryad: File 3). We did not assign a sampling fraction to genera of Icoisoptera because many are not monophyletic (Bourguignon *et al*., 2017; Buček *et al*., 2022b; Wang *et al*., 2023). The sampling fraction was assigned based on information available in Krishna *et al*. (2013) and the Termite Database (https://termitologia.net/; last accessed on 30/06/2024; (Constantino, 2024). We defined the sampling fraction as the number of species of a given clade present in our trees divided by the recorded species diversity of that clade. For example, our trees included 222 species of Nasutitermitinae, a subfamily currently represented by 604 described species (Krishna *et al*., 2013), so the sampling fraction for Nasutitermitinae was set to 0.36. For each of the four trees, the analyses were performed twice, once without the sampling fractions assigned to each clade (global analyses) and once with assigned sampling fractions (custom analyses; see Dryad: File 3).

Priors and parameters of the Markov chain Monte Carlo (MCMC) runs were estimated using the *setBAMMpriors* function implemented in the BAMMtools package. For each analysis, we performed four replicate MCMC runs of 50 million generations and sampled the parameters every 5,000 generations. We set the expected number of shifts to 50 to ensure no potential shifts were overlooked due to a small initial prior value, as recommended in the BAMM manual. Additionally, we set the *minCladeSizeForShift* to 5, which restricts rate-change events to internal branches with at least five descendant tips. This approach reduces computing resource usage and improves convergence. BAMM was run with a *segLength* setting of 0.02, splitting branches into fragments of 0.2% of the total tree height. This setting discretizes branches into intervals of approximately 2.54 Ma, ensuring satisfactory chain mixing and Effective Sample Size (ESS) values. For each 0.2% segment, a constant rate is assumed, enhancing computational efficiency. ESS values were monitored using Tracer v1.7.2 (Rambaut *et al*., 2018). The outputs from the diversification analyses performed with BAMM were processed with the R package BAMMtools to evaluate the shifts in diversification regimes across the termite tree of life and to visualize the evolutionary rates dynamics through time. All scripts, inputs, and outputs are available in Dryad (File 4).

In addition, we employed RPANDA v2.3 (Morlon *et al*., 2016) to estimate diversification rates over time and corroborate our BAMM results. We fitted six birth-death models accounting for the global sampling fraction of our phylogeny (0.39), with rates varying exponentially with time. The six models were a pure-birth constant rate model (BCST), a birth-death constant rate model (BCSTDCST), a speciation varying through time without extinction model (BTimeVar), a speciation varying through time with constant extinction rate model (BTimeVarDCST), a constant speciation rate with extinction varying through time model (BCSTDTimeVar), and a model with both speciation and extinction varying through time (BTimeVarDTimeVar). In the time-dependent models, *λ*, *μ*, or both vary as a continuous function of time *t* (Morlon *et al*., 2011), where *t* progresses from the present to the past. We assumed that these functions vary exponentially with *λ*(t) = *λ*_0_*e^αt^ and/or *μ*(t) = *μ*_0_*e^βt^, where *λ*_0_ and *μ*_0_ are the speciation and extinction rates at present, and α and β are the correlation coefficient specific to *λ* and *μ*. These coefficients measure the sign and pace, indicating how speciation and extinction rates have changed over time. Positive α and β suggest a slowdown in speciation and extinction processes towards the present, while negative α and β imply an acceleration of speciation and extinction processes. All scripts, inputs, and outputs are available in Dryad (File 5).

Finally, we conducted diversification analyses on our termite time-calibrated trees using the episodic birth-death model implemented in TreePar v3.3 (Stadler, 2011) to infer variations in speciation and extinction rates over time. TreePar implements a maximum-likelihood method that allows for rates to change at specific points in time, making it suitable for detecting rapid shifts in speciation and extinction rates in response to major events in Earth’s evolutionary history, such as drastic climate changes or gymnosperm-angiosperm turnover. We used the “*bd.shifts.optim*” function to estimate changes in speciation and extinction rates and their timing in the termite tree of life. We assessed potential diversification shifts every million years, allowing for up to ten shifts along the tree, including periods of declining diversity (negative diversification rates). To account for the incomplete taxon sampling, we applied an analytical correction using a global sampling fraction of 0.39, as described above. We fitted 11 diversification models differing in the number of diversification shifts (from zero to ten shifts) to our termite time-calibrated trees. The best-fit model was identified using the corrected Akaike Information Criterion (AICc). All scripts, inputs, and outputs are available in Dryad (File 6).

The models fitted with RPANDA and TreePar were compared using AICc and ΔAICc. The model with the lowest AICc is generally considered the best fit. Two recent shifts (estimated at 3 and 2 Ma) were evidenced in TreePar analyses for all four trees but were considered artefactual as they likely arose from our OTU-based tree-trimming approach, and models containing only these shifts were not identified as best fit by AICc (Supplementary Data 3). TreePar, RPANDA, and BAMM estimate speciation and extinction rates using different approaches. These methods were employed to explore various patterns of diversification. Since TreePar and RPANDA analyses rely on a model comparison framework (i.e., selecting the best-fit model from a set of predefined models), the results from these diversification analyses should be viewed as complementary to the results obtained with BAMM rather than as a cross-validation procedure.

### Addressing the identifiability issue

Macroevolutionary models of species diversification using speciation and extinction have faced criticisms regarding identifiability as an infinite number of models with different rates of speciation *λ* and extinction *μ* are equally likely (Louca & Pennell, 2020). In other words, rates of speciation *λ* and extinction *μ* cannot be unambiguously inferred from phylogenetic trees (Pagel, 2020), a problem that has been known since the first use of birth-death models to predict speciation and extinction rates from phylogenetic trees (Kubo & Iwasa, 1995). The unidentifiability of rates of speciation *λ* and extinction *μ*, which is characterized by identical likelihood for multiple values of this pair of parameters, affects all methods widely used for studying diversification, including those employed in this study. The issue of identifiability leads to the necessity of careful interpretation of the results (Helmstetter *et al*., 2022; Morlon *et al*., 2022) and implementation of a mitigation plan. Our macroevolutionary analyses were performed following previous recommendations (Rabosky *et al*., 2017; Rabosky, 2018; Morlon *et al*., 2022). First, we implemented priors within a Bayesian framework and penalized model complexity. Second, we conducted BAMM analyses on the most comprehensive termite tree of life reconstructed to date, focusing on testing hypotheses from the literature —*e.g.*, whether the diversification of crown Termitidae halted the diversification of Kalotermitidae; whether termite diversification coincides with periods of major climatic changes that affected tropical and subtropical ecosystems. Third, we explored the robustness of our BAMM results with complementary analyses using alternative birth-death models and different assumptions regarding the speciation and extinction processes. Fourth, while we estimated the rates of speciation *λ* and extinction *μ*, the conclusions of this paper primarily rely upon diversification rates *r*, which are not affected by the problem of identifiability described by Louca & Pennell (2020).

### Potential mechanisms of termite diversification

We used a birth-death model to investigate the potential impact of major past events in Earth’s evolutionary history on diversification rates (Condamine *et al*., 2013, 2019). We tested for correlations between past environmental changes and rates of speciation *λ* and extinction *μ* with the “*fit_env*” function implemented in RPANDA 2.3 (Morlon *et al*., 2016). This approach uses time-dependent diversification models and allows *λ* and *μ* to depend on time and external variables that vary over time. It assumes that clades evolve under a birth-death process, where *λ* and *μ* can vary with time and with one or more environmental variables.

We considered six environmental variables. First, we used two biotic variables, angiosperm (*A*) and gymnosperm (*G*) relative diversity, using available data from Silvestro *et al*. (2015) and Lehtonen *et al*. (2017). Second, we used four environmental abiotic variables: absolute temperatures (*T*), atmospheric concentrations in O2 (*O*) or CO2 (*C*), and continental fragmentation (*F*). We used the dataset of absolute temperatures compiled by Condamine *et al*. (2019) to model temperature variations. These temperature data did not reflect local or seasonal fluctuations but rather planetary-scale climatic trends, which may have resulted in temporally coordinated diversification changes across multiple clades. Changes in atmospheric O2 concentrations were modeled using the dataset of Lehtonen *et al*. (2017). Changes in CO2 concentrations during the Cretaceous-Cenozoic interval were modeled from the dataset compiled and refined by Foster *et al*. (2017). Finally, we used the dataset of Zaffos *et al*., (2017) to examine the effect of continental fragmentation on termite diversification. Given that environmental variables are often measured in discrete intervals, we used a smoothing function to model diversification rates in continuous time. We used the *pspline* v1.0-20 R package to generate datasets with 1-million-year intervals spanning from the Cretaceous to the present. All inputs used for these analyses are available in Dryad (File 5).

For the six considered environmental (E) variables, we used the same exponential dependencies as above when fitting diversification rates to time, that is: *λ*(E[t]) = *λ*_0_*e^αE[t]^ and *μ*(E[t]) = *μ*_0_*e^βE[t]^, with *E*[*t*]: *A*[*t*], *G*[*t*], *T*[*t*], *O*[*t*], *C*[*t*], or *F*[*t*]). In these cases, *λ*_0_ and *μ*_0_ are the expected speciation and extinction rates under an *E*[*t*] of 0, and α and β measure the sign and strength of the *E*[*t*] dependence. Hence, a positive α and β suggest that speciation and extinction rates are higher under high *E*[*t*] periods, while a negative α and β suggest that speciation and extinction rates are higher under low *E*[*t*] periods. We fitted and compared diversification models with constant rates (CST), time-dependent rates (TimeVar), temperature-dependent rates (TempVar), angiosperm-diversity-dependent rates (AngioVar), and gymnosperm- diversity-dependent rates (GymnoVar).

### Biogeographic stochastic mapping

We performed biogeographic analyses with R v4.2.3 using the package BioGeoBEARS v1.1.3 (Matzke, 2013). Ancestral area reconstruction was performed under the Dispersal-Extinction-Cladogenesis (DEC) model (Ree & Smith, 2008). We selected the DEC model without the *j* parameter and without time stratification because the DEC+*j* model generally predicts null or exceedingly low extinction rates and favors jump dispersals over widespread ancestral ranges, which is not suitable for reconstructing the history of old groups (Sanmartín & Meseguer, 2016; Rolland & Condamine, 2019). Reconstructions were carried out on the OTU trees using the realm of one randomly selected sample per tip. This approach reduces the influence of recent among-areas dispersal events and avoids inflating rates due to human-facilitated dispersals.

We considered six biogeographic realms adapted from Holt *et al*. (2013): Afrotropical, Australian, Neotropical, Nearctic, Oriental + Oceanian, and Palearctic + Sino-Japanese + Saharo-Arabian. We limited the number of areas as the DEC model is unable to handle a high number of regions in the geographic range evolution analyses (Antonelli *et al*., 2018). The DEC reconstruction was used as input to carry out 100 biogeographical stochastic mapping (BSM) using the ‘runBSM’ function (Dupin *et al*., 2017).

## Data Availability

Both the Supplementary Data and the Dryad Files listed in this manuscript are available to referees and will be made publicly available upon acceptance. All source data necessary to reproduce this study are available from the Dryad Digital Repository at: XXX. Samples used in this study are described in Supplementary Data 1. The UCE data from prior publications used in this study are available in the Dryad Digital Repository at: https://doi.org/10.5061/dryad.x0k6djhn0 (Hellemans *et al*., 2022b) (file: “Supplementary_Data_6_uces_TER_UCE_DB_CONTRIB_1.fasta”); https://doi.org/10.5061/dryad.5mkkwh77v (Buček *et al*., 2022a) (file: “TER_UCE_DB_CONTRIB_2.fasta.zip”); https://doi.org/10.5061/dryad.tmpg4f53w (Arora *et al*., 2023a) (file: “TER_UCE_DB_CONTRIB_3.tar.gz”); https://doi.org/10.5061/dryad.02v6wwqbm (Hellemans *et al*., 2024b) (file: “TER_UCE_DB_CONTRIB_4.fasta.gz”). The UCE dataset produced in this study is available on the Dryad Digital Repository (File 1), available at: XXX. All UCEs are centralized in the Termite UCE database, available at: https://github.com/sihellem/TER-UCE-DB/. Newly generated mitochondrial genomes are available on GenBank (see accession numbers in Supplementary Data 1). Additionally, we provide a curated termite mitogenome database encompassing all published records, available at: https://github.com/sihellem/TER-MT-DB/.

## Code Availability

No new custom code is published with this article. The Termite UCE database and associated codes are available at: https://github.com/sihellem/TER-UCE-DB/. The Termite Mitogenome database is available at: https://github.com/sihellem/TER-MT-DB/.

## Author Contributions Statement

SH, MW, and TB conceptualized the experiments. SH, MMR, JB, TFC, YR, RHS, and TB collected the samples. SH and TB supervised the project. MW performed lab experiments, generated the raw data, performed metagenome assemblies, identified and submitted the mitogenomes to GenBank. SH extracted the nuclear phylogenetic markers and performed the phylogenetic and biogeographic analyses. CJ performed the diversification analyses. SH, CJ, and TB wrote the original draft manuscript. SH prepared the display items. All authors (SH, MW, CJ, MMR, JB, TFC, FL, FLC, YR, EMC, RHS, and TB) edited and accepted the final version of this manuscript.

## Acknowledgments

We thank the Sequencing Section (SQC) and the Scientific Computing & Data Analysis Section (SCDA) of OIST for assistance with sequencing and providing access to the OIST computing cluster, respectively. This work was supported by subsidiary funding from OIST. SH was supported by the Japan Society for the Promotion of Science (JSPS) through a postdoctoral fellowship for foreign researchers (19F19819). TFC was funded by a grant from the São Paulo Research Foundation (FAPESP, #2020/06041-4). FLC benefited from an “Investissements d’Avenir” program managed by the Agence Nationale de la Recherche (CEBA, ref. ANR-10- LABX-25-01).

---

**Supplementary** Figure 1. Biogeographical reconstruction mapped on the UCEs_3CP- using the DEC model (without *j* parameter), and 100 biogeographic stochastic mapping (BSM) with BioGeoBEARS. We considered six biogeographic realms adapted from Holt *et al*. (2013): Afrotropical (green), Australian (dark pink), Neotropical (dark blue), Nearctic (light blue), Oriental + Oceanian (yellow), and Palearctic + Sino-Japanese + Saharo-Arabian (light violet). Operational Taxonomic Units (OTUs) denomination can be found in Supplementary Data 1.

