## Supplementary figures and images for "Termites became the dominant decomposers of the tropics after two diversification pulses"

### Supplementary Figure 1

**DEC model**

**ancstates: global optim, 6 areas max. d=0.0011; e=8e−04; j=0; LnL=−855.24**

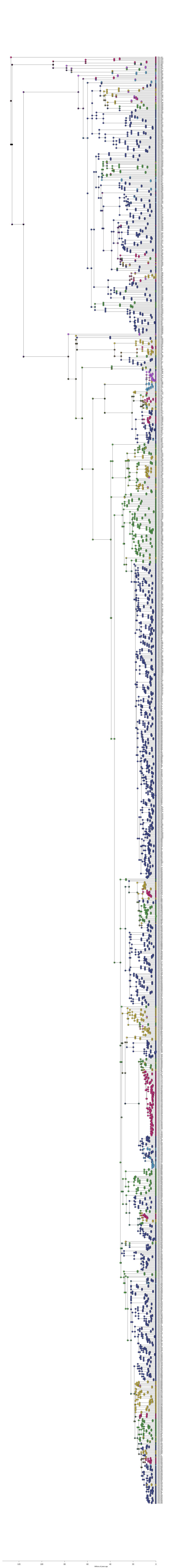
